# High-throughput profiling of natural acquired humoral immune response to a large panel of known and novel full length ectodomain *P. falciparum* merozoite vaccine candidates under reduced malaria transmission

**DOI:** 10.1101/2022.02.15.479108

**Authors:** Duncan Ndegwa Ndungu, James Tuju, Emily Chepsat, Rinter Mwai, Kennedy Mwai, Lydia Nyamako, Moses Mosobo, Awa B. Deme, Baba Dieye, Ibrahima Mbaye Ndiaye, Mouhamad Sy, Mamadou Alpha Diallo, Younous Diedhiou, Amadou Moctar Mbaye, Dyann Wirth, Daouda Ndiaye, Faith Osier, Amy K. Bei

## Abstract

Despite recent progress in the fight against malaria, it still remains a global health challenge necessitating development of intervention strategies. However, the search for malaria vaccine(s) has so far been very challenging. Multiple targets have been tested and so far only a few show promise with one having been endorsed by the WHO. In this study we explore the development of immunity in a low transmission setting, with very few documented re-infections, in order to understand the kinetics of the development and waning of immunity to current and novel blood-stage vaccine candidate antigens. To do this we performed a high-throughput measurement of natural acquired immunity against *P. falciparum* antigens utilizing a well-established micro-array platform based on the mammalian protein expression system. This large panel of known and novel recombinant full length ectodomain *P. falciparum* merozoite vaccine candidates were differently recognized by the immune system. Based on the overal spread of the data, some of these antigens induced the acquisition of high levels (1^st^ tertile) of antibodies, among which included novel antigens such as PF3D7_1025300, PF3D7_1105800, PF3D7_1334400, PF3D7_0911300, PF3D7_1252300, PF3D7_1460600, PF3D7_1453100, PF3D7_0831400 and some induced low levels of antibodies (3rd tertile) while others induced moderate levels (4th tertile). In this longitudinal cohort with low level of malaria endemicity, acquisition of humoral immunity to these full length ectodomains *P. falciparum* antigens demonstrate different dynamics over-time, whereby it was either not acquired or if it was acquired it was either maintained or lost at different rates. These various identified novel antigens are potentially ideal candidates to be prioritized for further functional and or serological studies.

## INTRODUCTION

Malaria is a disease caused by parasites of the species *Plasmodium.* Among the five *Plasmodium* species that can cause malaria in humans, the majority of clinical cases are caused by *P. falciparum* and *P. vivax* with the former mainly being the major cause of the malaria cases in Africa. Despite recent progress in the fight against malaria, progress has plateaued, and malaria remains a global health challenge (World Malaria Report 2019). As of 2019, malaria caused over 200 million cases and more than 400,000 deaths (World Malaria Report 2020), most of which were African children, therefore necessitating rapid development of intervention strategies. The complex life-cycle of the human malaria parasites present multiple potential targets for these intervention strategies. These include targeting the liver, blood and mosquito stages of the parasite as well as the mosquito vector to prevent disease and the transmission of the parasite from one infected individual to another. However, this vector of the malaria parasite continues to develop resistance to pesticides while the complex biology of this parasite has so far been an obstacle in the development of a permanent cure for complete elimination of this disease. The malaria parasite continues to develop resistance to anti-malarial drugs. Despite huge efforts to develop malaria vaccine(s) with multiple targets having been tested (Reviewed in (Barry and Arnott 2014; Duffy and Patrick Gorres 2020)) only a few show promise: the malaria pre-erythrocytic vaccine RTS,S which was recently approved for use by the WHO, and its improved version R21 (Datoo et al. 2021).

In malaria endemic countries, repeated exposure to malaria infection from childhood to adulthood results in natural acquired immunity to malaria disease, but does not result in sterilizing immunity from infection and antibodies purified from serum obtained from malaria immune adults has been used to treat non-immune malaria patients (Cohen, McGREGOR, and Carrington 1961; Sabchareon et al., 1991), providing compelling evidence of the importance of antibodies in the immune response to malaria that can control parasitemia (Cohen, McGREGOR, and Carrington 1961). Protective immunity to malaria is associated with the recognition of a combination of antigens (Proietti et al. 2020; Bustamante et al. 2017; Daou et al. 2015; Osier et al. 2014; Yman et al. 2020) and development of antibodies to *P. falciparum* antigens play a role in protection from malaria disease (Cohen, McGREGOR, and Carrington 1961). But identifying the dominant protective candidates among these potential 5400 (Gardner et al. 2002) proteins in the *P. falciparum* proteome has remained a challenge. In-spite of this and even though during pre-genomic era the number of candidates of interest was always few, various studies managed to identify malarial vaccine candidates (Sirima et al. 2016; Chitnis et al. 2015; Ogutu et al. 2009; Sheehy et al. 2011; 2012; Bélard et al. 2011; Thera et al. 2011; Srinivasan et al. 2017; Duncan et al. 2011; Ellis et al. 2012; Biswas et al. 2014; Koram et al. 2016; El Sahly et al. 2010; Lawrence et al. 2000; Genton et al. 2002, 2003; Salamanca et al. 2019; Beeson et al. 2016). But because the focus was on antigens that are generally immunodominant and didn’t represent the global parasite population, these candidates didn’t succeed mainly due to antigenic diversity and strain specific immunity (Ouattara et al. 2013; Genton et al. 2002, 2003; Thera et al. 2011; Barry et al. 2009; E. et al. 2012; Takala et al. 2009; Barry and Arnott 2014; Bailey et al. 2020). Now with the completion of the sequencing of the malaria parasite genome (Gardner et al. 2002) coupled with the mining of genomics, transcriptomics or proteomic data, its now possible to screen the whole proteome of the malaria parasite to prioritise vaccine candidates (reviewed in (Tuju et al. 2017; Proietti and Doolan 2015)). Indeed, using microarray, various studies have probed the malarial proteome for antigens that could be used for vaccine candidate development, malaria diagnosis, malaria surveillance or to generally understand the immunology of the malaria parasite (Bailey et al. 2020; Camponovo et al. 2020; Liang and Felgner 2015; Yman et al. 2020; Proietti et al. 2020; Zhou et al. 2019; Obiero et al. 2019; Kobayashi et al. 2019; van den Hoogen et al. 2019; Jaenisch et al. 2019; Taghavian et al. 2018; Kamuyu et al. 2018; Uplekar et al. 2017; Helb et al. 2015; Dent et al. 2015; Finney et al. 2014; Baum et al. 2013; Fan et al. 2013; Crompton et al. 2010; Doolan et al. 2008; Proietti et al. 2020; Kobayashi et al. 2019).

As opposed to the whole malaria parasite antigen which induces rapid and long-lasting antibody, humoral immunity to single antigens develops slowly (Corran et al. 2007). This humoral immune responses to individual malaria antigens display different dynamics of acquisition and decay depending on the intensity of transmission (Ondigo et al., 2014). This immunity can persist over months or years (Corran et al. 2007) followed with decrease over time in an antigen specific manner, making it possible to select biomakers that can be used to predict recent and distant malaria exposure (Helb et al. 2015). As malaria transmission goes down as we are achieving the goal of regional malaria elimination, more malaria endemic settings will look like the situation in Senegal – low sporadic transmission. Thus this study can teach us about what happens to immunity in a low transmission setting.

Previous malaria seroepidemiology utilised cell free protein expression systems such as *E. coli* and wheat germ to express *Plasmodium* antigens. But with recent technological advancements the mammalian protein expression system has now been adapted making it possible to overcome some of the previous challenges such as incorrect folding post-translation modification and potential missing of immunodominant epitopes (Crosnier et al. 2013; Zenonos, Rayner, and Wright 2014) which could therefore interfere with epitope integrity, antigen presentation and consequently their recognition by the host immune system. In this study our main objective is to study natural immune responses in a low-transmission setting to determine the kinetics of antibody acquisition and decline, without frequent re-exposure. To achieve this we performed a high-throughput measurement of natural acquired immunity against *P. falciparum,* utilizing the well-established Kilchip micro-array plat-form (Kamuyu et al. 2018), which is a multiplex large panel of mostly full length protein ectodomains (about 70% in total) of novel and known malaria vaccine candidates that were selected based on the *P. falciparum* merozoite surface localized, secreted and invasion proteome, and then expressed mainly using the mammalian protein expression system. We show that natural immunity to this large panel of known and novel recombinant full length ectodomain *P. falciparum* merozoite vaccine candidates is acquired, maintained, and lost with unique kinetics to what has been described in other settings. Understanding the unique kinetics of antibody acquisition, maintenance, and reversion can inform frameworks for development of the next generation of malaria vaccine candidates.

## RESULTS

### Study Sites and Patient Demographics

Study participants consisted of a total of 70 male patients with a median age of 11 (5-16) years, median parasitemia on the day of recruitment of 0.43(0.02-4.89)% and with a mean of 1 (0-5) number of past malaria infections (Table 1), described previously (Sy et al. 2020). These participants were recruited from six neighborhoods of the Thiès region of Senegal. These neighborhoods include Diakhao, Nasrou, Takhikao, Thially, Cite Senghor and Escale. The participants were followed for a total period of two years with scheduled follow-up and sample collection done on Days 0, 14, 28, 92, 183, 365, 543 and 730 (Supplementary Table 1). Only 4 participants had malaria re-infections over the 2 year follow-up period, defined based on symptoms and confirmed with RDT and microscopy. These unscheduled visits occurred on days 326, 374 and 397 (Supplementary Table 1).

**Table 1.**

|  | Diakhao | Nasrou | Takhikao | Thially | Cite Senghor | Escale |
| --- | --- | --- | --- | --- | --- | --- |
| **Number of participants** | 47 | 11 | 5 | 5 | 1 | 1 |
| **Median age** | 10(5-15) | 11(8-16) | 11 | 12(8-14) | 13 | 10 |
| **Gender** | M | M | M | M | M | M |
| **Median percent parasitemia on Day0** | 0.45(0.03-4.27) | 0.43(0.03-3.01) | 0.015(0.06-2.28) | 1.01(0.02-4.89) | NA | 0.03 |
| **Median number of past malaria infections** | 1(0-5) | 0(0-1) | 0(0-2) | 0 | 0 | 2 |

### A large panel of known and novel recombinant full length ectodomain *P. falciparum* merozoite vaccine candidates are recognized by the humoral immune system

To determine the kinetics of antibody acquisition and decline, without frequent re-exposure we utilised the well-established Kilchip micro-array platform (Kamuyu et al. 2018) to perform a high-throughput measurement of antibody recognition of a large panel of mostly full-length protein ectodomains of novel and known malaria vaccine candidates expressed mainly using the mammalian protein expression system against serum from the above cohort. We focused on full length ectodomains selected from the merozoite proteome (Table 2). These selected antigens are localized on different regions of the merozoite (Figure 1) which include: 1. Secreted proteins: PF3D7_0323400 (RIPR), PF3D7_0207900 (SERA2), PF3D7_1334400 (MSRP4),PF3D7_0902800 (SERA9), PF3D7_0911300 (MSP3.6), PF3D7_0207700 (SERA4), PF3D7_0207400 (SERA7), PF3D7_0207800 (SERA3), PF3D7_1017100 (100166), PF3D7_0911300 (MSP3.5), PF3D7_0104200 (PF210C), PF3D7_0424400 (B29), PF3D7_0424400 (B27), PF3D7_0424400 (B30), PF3D7_0424400 (B31) (Zhu et al. 2017; Zenonos, Rayner, and Wright 2014; van Ooij et al. 2013); 2. Surface or GPI-Anchored proteins: PF3D7_0502400 (MSP8), PF3D7_0707300 (RAMA), PF3D7_0102700 (0135W), PF3D7_0206800 (MSP2), PF3D7_0612700 (P12), PF3D7_0930300 (MSP1), PF3D7_1420700 (P113), PF3D7_0620400 (MSP10), PF3D7_0207000 (MSP4), PF3D7_0508000 (P38), PF3D7_1136200 (1136200), PF3D7_0206900 (MSP5), PF3D7_1364100 (PF92), PF3D7_1401600 (1401600) (Crosnier et al. 2013; Zenonos, Rayner, and Wright 2014; Ntumngia et al. 2004); 3. Microneme proteins: PF3D7_1133400 (AMA1), PF3D7_0731500 (EBA175), PF3D7_1301600 (EBA140), PF3D7_0102500 (EBA181), PF3D7_0405900 (ASP), PF3D7_1028700 (MTRAMP), PF3D7_0828800 (GAMA) (13); 4. Peripheral proteins: PF3D7_1035400 (MSP3), PF3D7_0404900 (P41), PF3D7_1036000 (MSP11), PF3D7_1035500 (MSP6), PF3D7_1228600 (MSP9), PF3D7_1334600 (MSRP3), PF3D7_1335100 (MSP7) (Crosnier et al. 2013); 5. Rhoptry proteins: PF3D7_0424100 (RH5), PF3D7_0402300 (RH1), PF3D7_0212600 (SPATR), PF3D7_0423400 (AARP), PF3D7_0905400 (RHOP3), PF3D7_0302200 (CLAG3.2) (Crosnier et al. 2013; Kalra et al. 2016; Holder et al. 1985); 6. Membrane proteins: PF3D7_1462300 (1462300), PF3D7_1460600 (1460600) (Morse et al. 2016; Poulin et al. 2013; Wang et al. 2020); 7. Parasitophorous vacuole membrane: PF3D7_1021800 (PFSEA), PF3D7_1033200 (ETRAMP) (Raj et al. 2014; Spielmann, Fergusen, and Beck 2003); 8. Dense granules: PTEX150 (Bullen et al. 2012) and others with no localization data: PF3D7_0606800 (606800), PF3D7_0616500 (TLP), PF3D7_1345100 (1345100), PF3D7_1453100 (145300), PF3D7_1105800 (1105200), PF3D7_0925900 (925900), PF3D7_0525800 (525800), PF3D7_0830500 (83500), PF3D7_1237900 (1237900), PF3D7_0206200 (206200), PF3D7_1229300 (1229300), PF3D7_1025300 (1025300), PF3D7_1252300 (1252300), PF3D7_0419700 (PF34), PF3D7_0831400 (831400), PF3D7_1334300 (MSRP5), PF3D7_1137300 (113700), PF3D7_0730800 (730800.2). These antigens were expressed using the mammalian protein expression system then they were printed on micro-array slides and detected as described in (Kamuyu et al. 2018) against a panel of whole serum donated by malaria infected children with low previous malaria infection history resident in a lowly malarial endemic region of Senegal (Table 1). The resulting reactivity measured as median fluorescent intensity (MFI) relative to mean MFI plus 2 standard deviations of non-immune control serum varied from strong to weak with an average of 0.666104 (range −0.88-2.34) with the respective antigens being lowly, averagely or highly recognized by the test serum (Figure 2A and 2B).

**Table 2.**

|  | **PlasmoDB ID** | **Gene Name** | **Abbreviation 1** | **Abbreviation 2** |
| --- | --- | --- | --- | --- |
| 1 | PF3D7_0502400 | merozoite surface protein 8 | MSP8 | 12MSP8 |
| 2 | PF3D7_0707300 | rhostry-associated membrane antigen | RAMA | RAMA |
| 3 | PF3D7_1133400 | apical membrane antigen 1 (AMA1) | AMA1 | AMA1 |
| 4 | PF3D7_1301600 | erythrocyte binding antigen-140 | EBA140 | 1EBA140 |
| 5 | PF3D7_0731500 | erythrocyte binding antigen-175 | EBA175 | 2EBA175 |
| 6 | PF3D7_0102500 | erythrocyte binding antigen-181 | EBA181 | 3EBA181 |
| 7 | PF3D7_0405900 | apical sushi protein | ASP | ASP |
| 8 | PF3D7_0828800 | GPI-anchored micronemal antigen | GAMA | GAMA |
| 9 | PF3D7_1028700 | merozoite TRAP-like protein | MTRAMP | MTRAMP |
| 10 | PF3D7_0606800 | VFT1 | 606800 | 606800 |
| 11 | PF3D7_1033200 | early transcribed membrane protein 10.2 | ETRAPM | ETRAPM |
| 12 | PF3D7_1021800 | schizont egress antigen-1 | PFSEA | 3PFSEA |
| 13 | PF3D7_0616500 | TRAP-like protein | TLP | TLP |
| 14 | PF3D7_1036000 | merozoite surface protein 11 | MSP11 | 3MSP11 |
| 15 | PF3D7_1035400 | merozoite surface protein 3 | MSP3 | 5MSP3 |
| 16 | PF3D7_1035500 | merozoite surface protein 6 | MSP6 | 10MSP6 |
| 17 | PF3D7_1228600 | merozoite surface protein 9 | MSP9 | 13MSP9 |
| 18 | PF3D7_0404900 | 6-cysteine protein P41 | P41 | P41 |
| 19 | PF3D7_1335100 | merozoite surface protein 7 | MSP7 | 11MSP7 |
| 20 | PF3D7_1334600 | MSP7-like protein | MSRP3 | 1MSRP3 |
| 21 | PF3D7_0423400 | apical asparagine-rich protein AARP | AARP | AARP |
| 22 | PF3D7_0905400 | high molecular weight rhostry protein 3 | RHOP3 | RHOP3 |
| 23 | PF3D7_0424100 | reticulocyte binding protein homologue 5 | RH5 | 2RH5 |
| 24 | PF3D7_0212600 | secreted protein with altered thrombospondin repeat domain | SPATR | SPATR |
| 25 | PF3D7_0402300 | reticulocyte binding protein homologue 1 | RH1 | 1RH1 |
| 26 | PF3D7_0911300 | cysteine repeat modular protein 1 | MSP3.5 | 6MSP3.5 |
| 27 | PF3D7_1334400 | MSP7-like protein | MSRP4 | 2MSRP4 |
| 28 | PF3D7_0104200 | StAR-related lipid transfer protein | PF210C | PF210C |
| 29 | PF3D7_0207900 | serine repeat antigen 2 | SERA2 | 1SERA2 |
| 30 | PF3D7_0207700 | serine repeat antigen 4 | SERA4 | 3SERA4 |
| 31 | PF3D7_0902800 | serine repeat antigen 9 | SERA9 | 5SERA9 |
| 32 | PF3D7_1017100 | rhostry neck protein 12 | 100166 | 100166 |
| 33 | PF3D7_0911300 | cysteine repeat modular protein 1 | MSP3.6 | 7MSP3.6 |
| 34 | PF3D7_0323400 | Rh5 interacting protein | RIPR | RIPR |
| 35 | PF3D7_0207800 | serine repeat antigen 3 | SERA3 | 2SERA3 |
| 36 | PF3D7_0207400 | serine repeat antigen 7 | SERA7 | 4SERA7 |
| 37 | PF3D7_0102700 | MaTrA | 0135W | 0135W |
| 38 | PF3D7_0620400 | merozoite surface protein 10 | MSP10 | 2MSP10 |
| 39 | PF3D7_0930300 | merozoite surface protein 1 | MSP1 | 1MSP1 |
| 40 | PF3D7_0206800 | merozoite surface protein 2 | MSP2 | 4MSP2 |
| 41 | PF3D7_0207000 | merozoite surface protein 4 | MSP4 | 8MSP4 |
| 42 | PF3D7_1420700 | surface protein P113 | P113 | P113 |
| 43 | PF3D7_1136200 | PF3D7_1136200 | 1136200 | 1136200 |
| 44 | PF3D7_0206900 | merozoite surface protein 5 | MSP5 | 9MSP5 |
| 45 | PF3D7_0508000 | 6-cysteine protein | P38 | P38 |
| 46 | PF3D7_0612700 | 6-cysteine protein P12 | P12 | P12 |
| 47 | PF3D7_1364100 | 6-cysteine protein P92 | PF92 | 2PF92 |
| 48 | PF3D7_1025300 | PF3D7_1025300 | 1025300 | 1025300 |
| 49 | PF3D7_1105800 | PF3D7_1105800 | 1105200 | 1105200 |
| 50 | PF3D7_1252300 | PF3D7_1252300 | 1252300 | 1252300 |
| 51 | PF3D7_0206200 | pantothenate transporter | 206200 | 206200 |
| 52 | PF3D7_0831400 | Plasmodium exported protein | 831400 | 831400 |
| 53 | PF3D7_0925900 | lipocalin(LCN), parasitophorous vacuolar protein 5(PV5) | 925900 | 925900 |
| 54 | PF3D7_1453100 | dynactin subunit 4 | 145300 | 145300 |
| 55 | PF3D7_1460600 | inner membrane complex sub-compartment protein 3 | 1460600 | 1460600 |
| 56 | PF3D7_0424400 | surface-associated interspersed protein 4.2 (SURFIN 4.2) | B29 | 20B29 |
| 57 | PF3D7_0419700 | apical merozoite protein | PF34 | 1PF34 |
| 58 | PF3D7_1436300 | translocon component PTEX150 | PTEX150 | PTEX150 |
| 59 | PF3D7_0525800 | inner membrane complex protein 1g (IMC1g) | 525800 | 525800 |
| 60 | PF3D7_0730800 | Plasmodium exported protein | 730800.2 | 730800.2 |
| 61 | PF3D7_0830500 | TryThrA | 83500 | 83500 |
| 62 | PF3D7_1137300 | CLPTM1 domain-containing protein | 113700 | 113700 |
| 63 | PF3D7_1229300 | PF3D7_1229300 | 1229300 | 1229300 |
| 64 | PF3D7_1237900 | PF3D7_1237900 | 1237900 | 1237900 |
| 65 | PF3D7_1345100 | Thioredoxin 2 | 1345100 | 1345100 |
| 66 | PF3D7_1401600 | Plasmodium exported protein (PHISTb) | 1401600 | 1401600 |
| 67 | PF3D7_1462300 | GTP-binding protein | 1462300 | 1462300 |
| 68 | PF3D7_0424400 | surface-associated interspersed protein 4.2 (SURFIN 4.2) | B27 | 18B27 |
| 69 | PF3D7_0424400 | surface-associated interspersed protein 4.2 (SURFIN 4.2) | B30 | 22B30 |
| 70 | PF3D7_0424400 | surface-associated interspersed protein 4.2 (SURFIN 4.2) | B31 | 23B31 |
| 71 | PF3D7_1334300 | MSP7-like protein | MSRP5 | 3MSRP5 |
| 72 | PF3D7_0302200 | cytoadherence linked asexual protein 3.2 | CLAG3.2 | CLAG3.2 |

**Figure 1:**
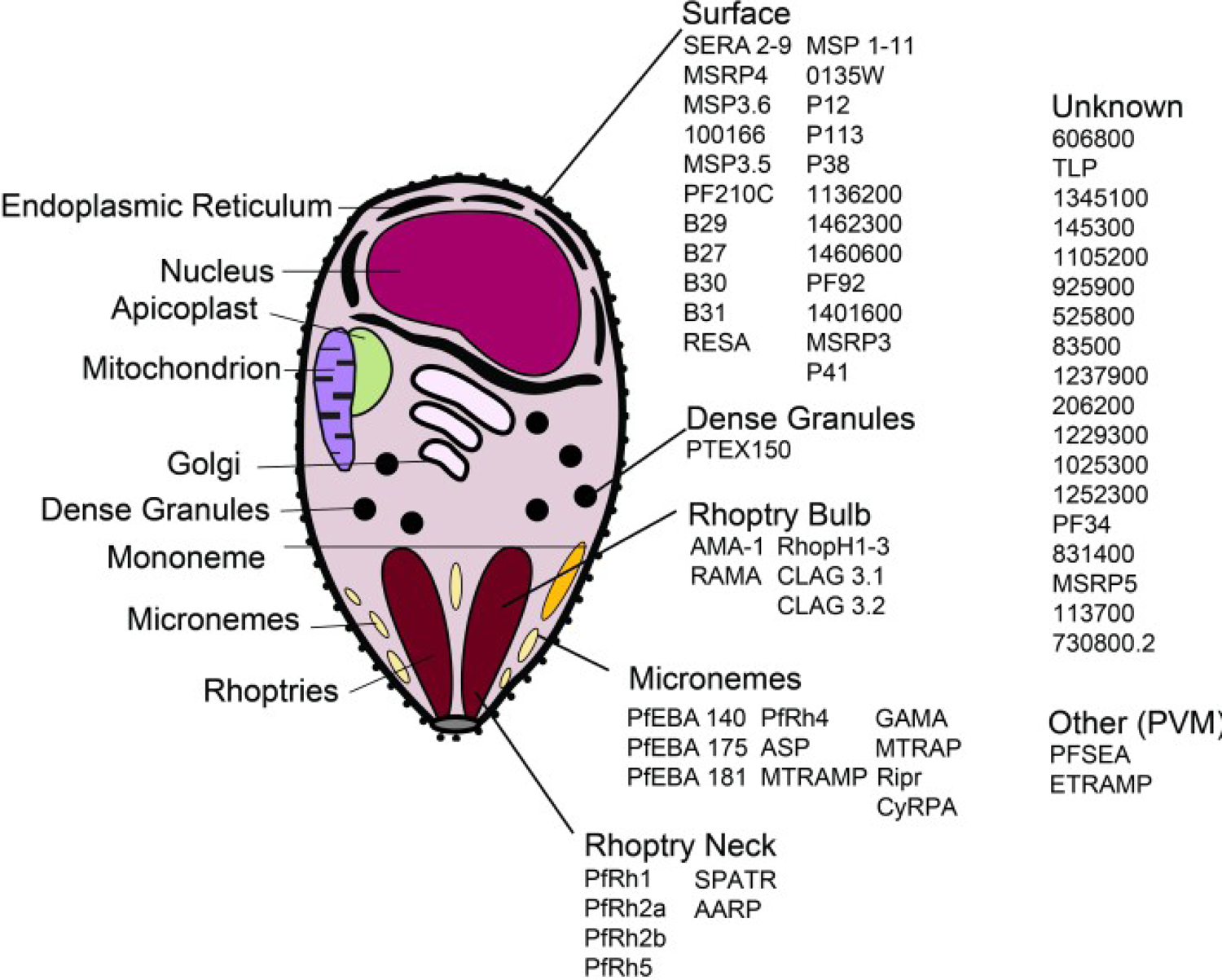
Localization of selected antigens on different regions of the merozoite. Selected antigens included those localised on the surface, micronemes, rhoptries, dense granules, parasitophorous vacuole and others of unknown localisation.

**Figure 2:**
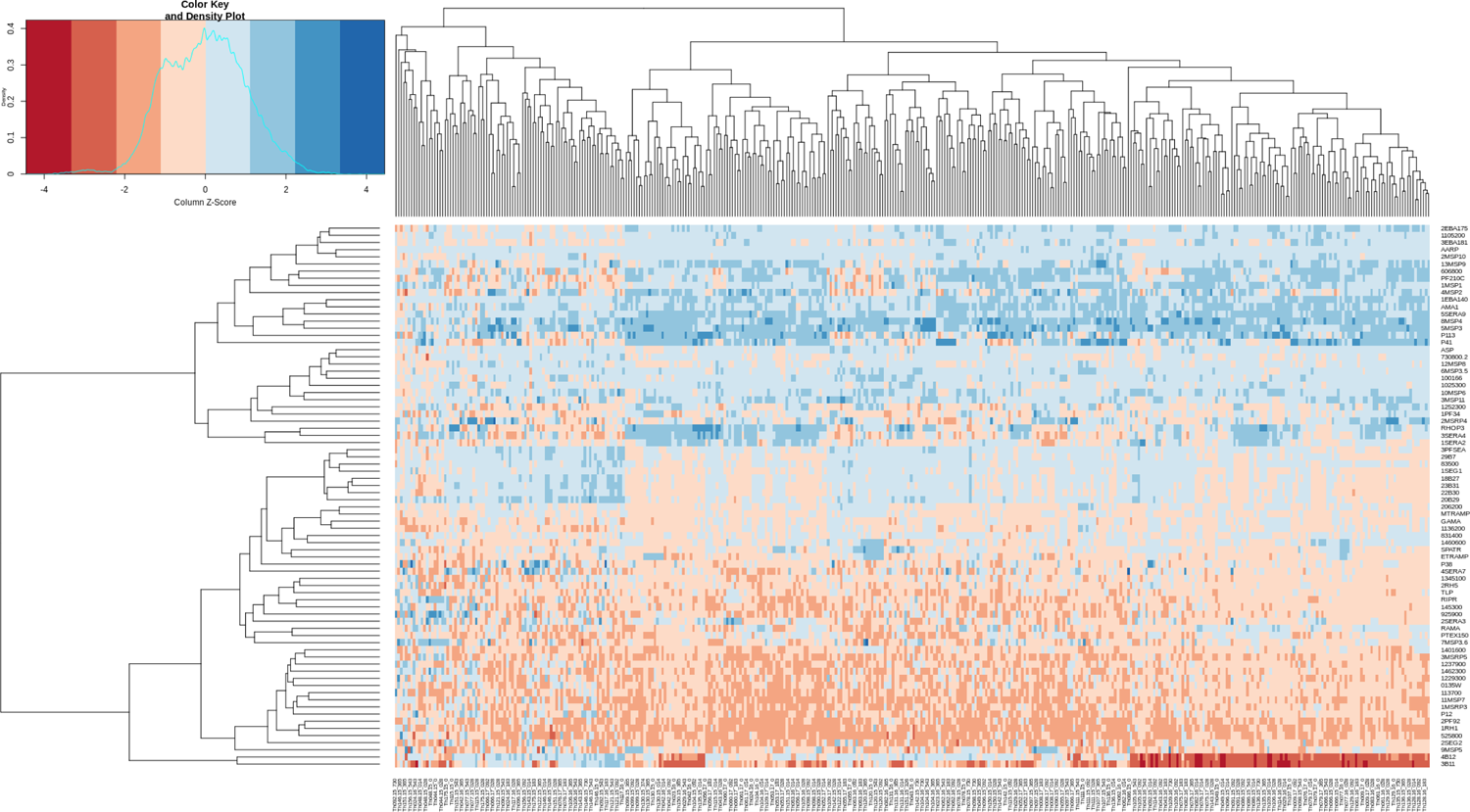
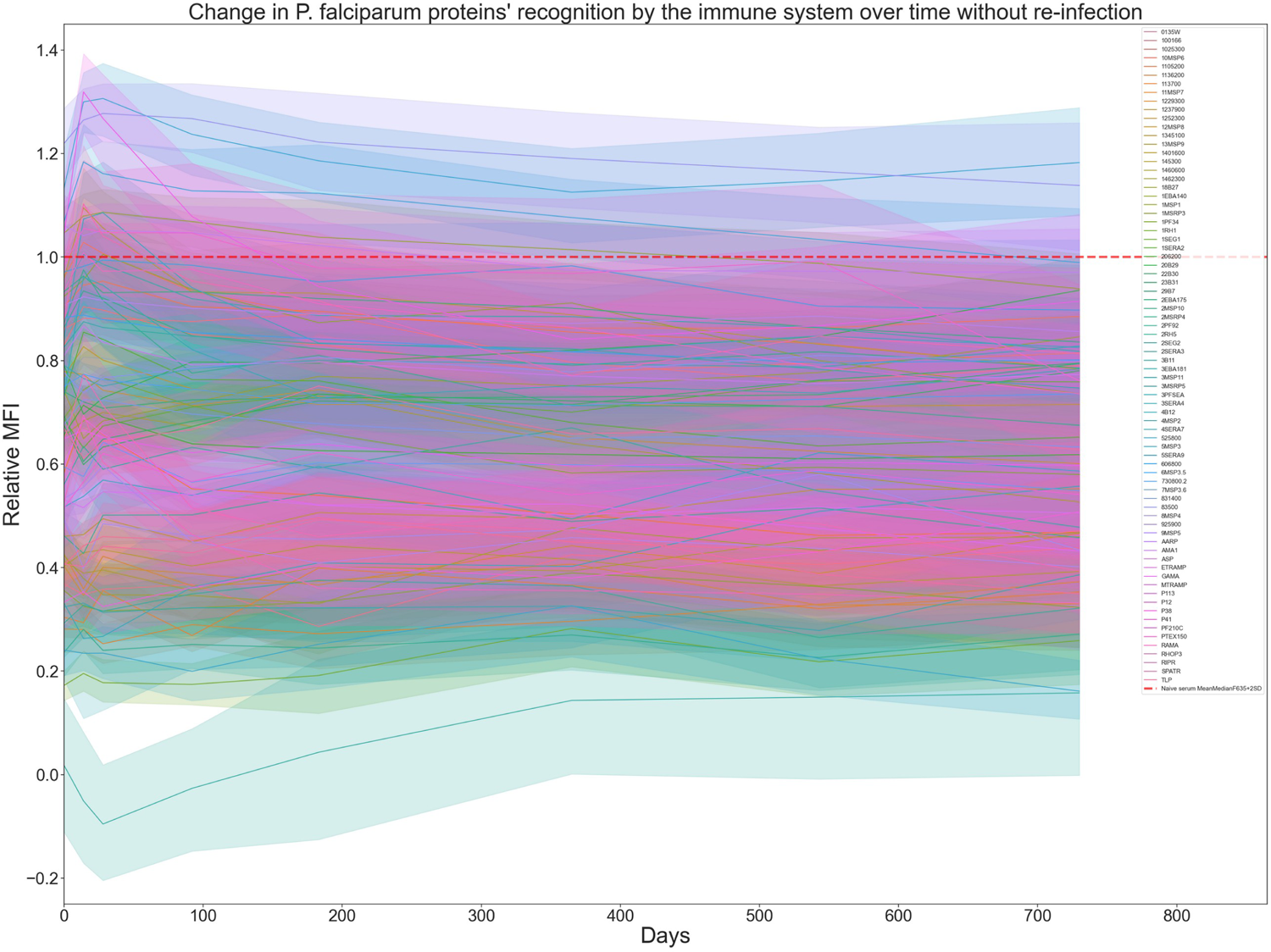
Serological response to merozoite antigens (known and novel vaccine candidates) in a lowly endemic region of Senegal (response and dynamics with time). A). Antibody reactivity against a large panel of known and novel P. falciparum merozoite vaccine candidates. A heat-map of normalized back-ground corrected median fluorescent intensity(MFI) relative to mean MFI plus 2 standard deviations of 8 non-immune control serum representing recognition of test protein (Vertical axis) by patient serum from the day of contact (horizontal axis) clustered using hierarchical clustering. Red represents low MFI while blue represents high MFI. B). Change in recognition of *P. falciparum* proteins over time of follow-up without re-infection. MFI of each test protein as recognized by test serum per each contact day (Y-axis) is plotted relative to mean MFI plus 2 standard deviations of non-immune control serum (red line) over the whole study period (X-axis).

### Full length ectodomains *P. falciparum* merozoite vaccine candidates induce three distinct patterns of antibodies acquisition

To measure antibody acquisition to *P. falciparum* antigens, the MFI of each test protein (Figure 2A) as recognized by test serum per each contact day was divided by mean MFI plus 2 standard deviations of non-immune control serum. Samples with values above 1 were considered as seropositive while samples with values below 1 were considered as seronegative. This indicates that some antigens are able to induce antibody acquisition to malaria infections while some are not. To measure the levels of these acquired antibody against *P. falciparum* antigens in the sampled population and in order to determine the most immunogenic antigen, the total number of sero-positive serum samples per each protein over the whole study period was determined. These values were then ranked from highest to lowest, divided into three groups based on the MFI tertiles and labeled respectively as 1 ^st^, 2^nd^ and 3^rd^ tertile (Figure 3A and supplementary table 2). This analysis was done in the absence of the individuals with re-infection, thus all data is from a single infection or boosting alone.

**Figure 3:**
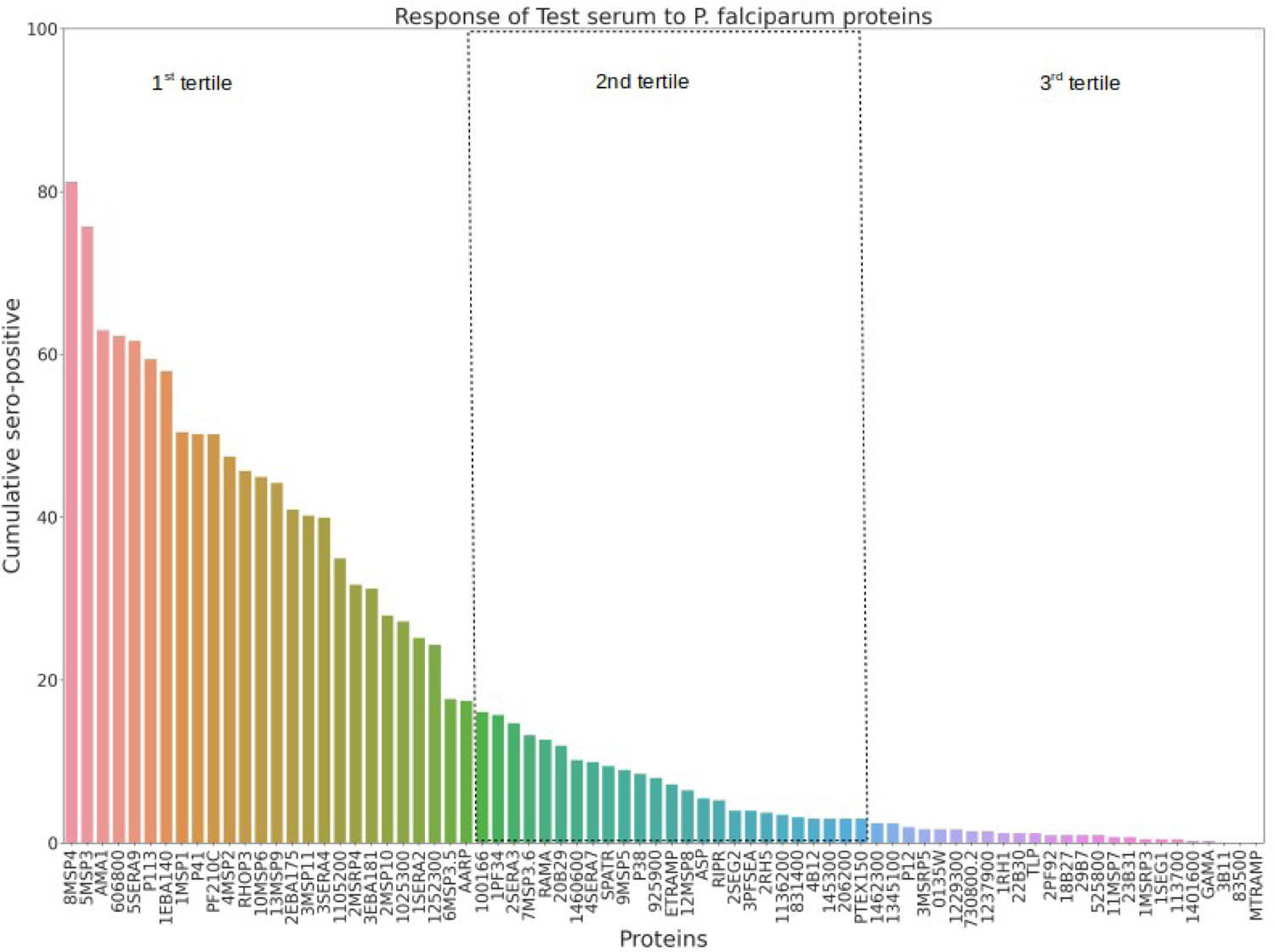
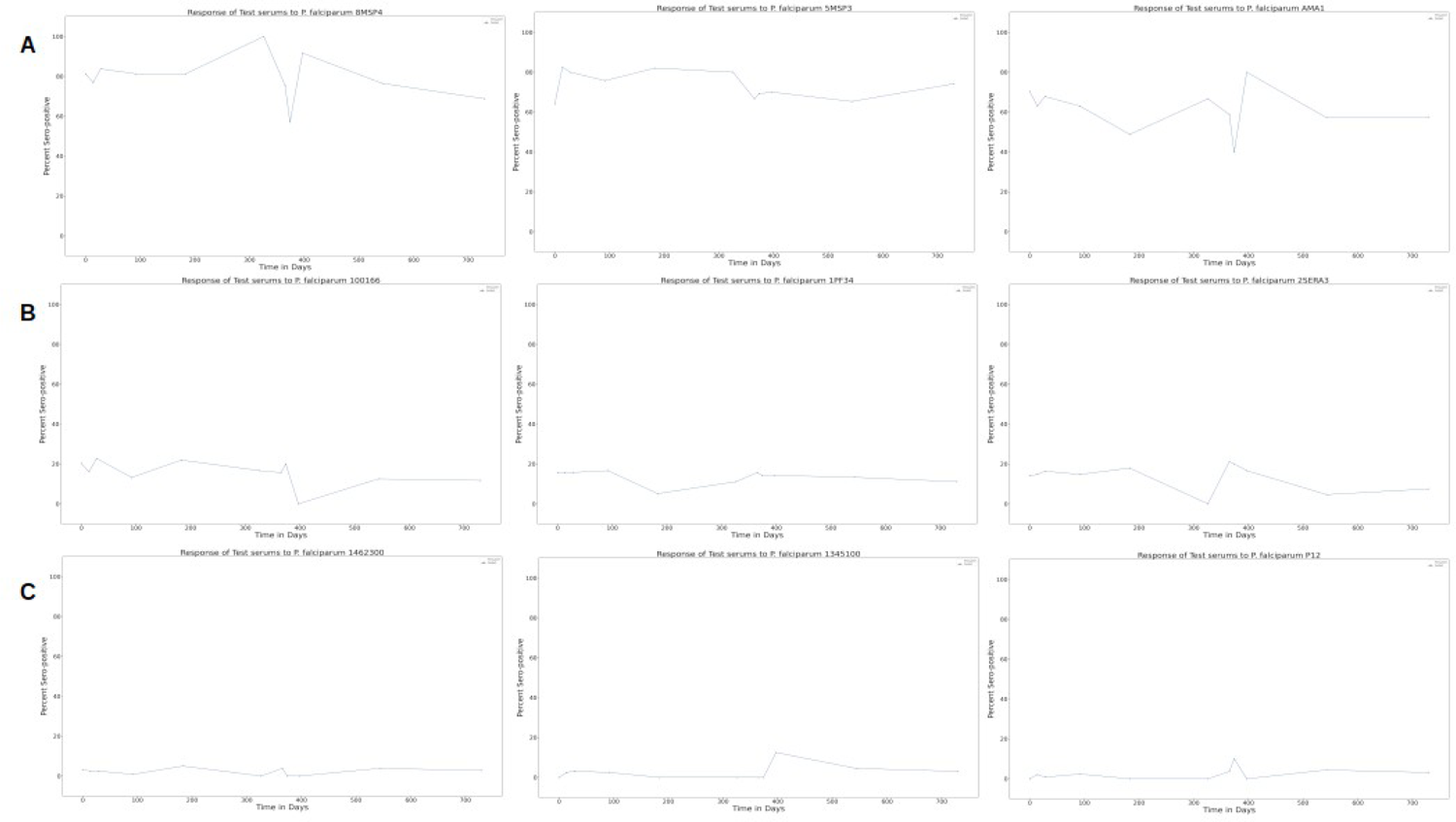
Antibody acquisition to a large panel of known and novel *P. falciparum* **merozoite full length ectodomains vaccine candidates. A). Levels of acquired antibody against *P. falciparum* antigens in the sampled population.** Y-axis is the total number of sero-positive serum samples per each protein on the X-axis over the whole study period. These values were then ranked from highest to smallest and divided into three groups. The proteins in the highest recognized category are indicated as 1^st^ tertile while the least recognized are indicated as 2^nd^ tertile. The moderately recognized proteins are indicated as 3^rd^ tertile. **B). Change in the levels of acquired antibodies against *P. falciparum* antigens**. Line plots of percent sero-positive test serums per each contact day (Y-axis) over the whole study period (X-axis) against representative test proteins from each of the three tertiles.

With a mean cumulative percent sero-positivity of 44.40% (range 16.82-81.25%) the proteins in the 1^st^ tertile were the most reactive, followed with those in the 2^nd^ tertile with a mean cumulative percent sero-positivity of 7.79%(range 3.00-16.13%) while those in the 3^rd^ tertile were least reactive with a mean cumulative percent sero-positivity of 1.06%(range 0.00-2.50%) (supplementary table 2). Antigens in the highest tertile included known dominant serodominant antigens (AMA-1, MSP-1, other MSPs, RHOP3, SERAs) any of which are merozoite surface antigens, in addition to previously un-explored targets (1105200, MSRP4, MSP3.5, 1252300, SERA2, 1025300). Proteins in the second tertile included invasion ligands (RIPR, RH5), components of a translocon (PTEX150), PVM components (PFSEA, ETRAMP), and previously unexplored targets (145300, 1460600, 831400, 206200, 925900, B29).

Proteins in the third tertile included previously unexplored targets (1229300, 1462300, 1237900, 730800.2, 113700, 525800, B30, 1345100, 83500, B31 and B27), redundant/dispensable antigens (P12, RH1, 1401600) and others (0135W, MSRP5, MSRP3, MTRAMP, PF92, MSP7, TLP, GAMA).

To measure change in the levels of these acquired antibodies in this cohort we determined the percentage of serum samples that were sero-positive per test proteins per each contact day during the study period. This was done separately using a) samples without malaria reinfection and b) samples without malaria re-infection and with malaria re-infection. The percent seropositive test serums per each contact day over the whole study period varied in an antigen specific manner that could be generalized into three groups based on the above observed tertiles (Figure 3B). Antigens falling under the 1st tertiles (Figure 3A) generally had the highest percent sero-positivity (Figure 3B and supplementary figure 1A) and antigens falling under the 3^rd^ tertiles (Figure 3A) generally had the lowest percent sero-positivity (Figure 3B and supplementary figure 1A) while those in the 2^nd^ tertile (Figure 3A) generally had moderate percent sero-positivity (Figure 3B and supplementary figure 1A). Confirmed re-infection with *P. falciparum* did not have a major influence on percent sero-positivity (supplementary figure 1B). Only in 28% of the antigens did boosting of percent sero-positivity occur (supplementary figure 1C). These antigens include 1RH1, 4MSP2, P38, 5MSP3, ETRAMP, 606800, SPATR, PF210C, 1025300, 1SERA2, AMA1, 1460600, 8MSP4, P113, 5SERA9, 4SERA, P41, 1PF34, 3MSP11, 3SERA4, RHOP3. The boosting in majority of these antigens occurred between day 300 and 400. In the remaining few, boosting had a different pattern. In 8MSP4 it occurred through-out the study period, in RHOP3 it occurred from the begining of the study period to day 400, in AMA1 it occurred between day 100 and around day 325, in ETRAMP P41 and P113 boosting occurred at multiple time-points between day 0 and day 400.

### Kinetics of naturally acquired antibodies to next generation malaria vaccine targets

Longevity of antibodies to *P. falciparum* antigen is antigen specific (Akpogheneta et al. 2008). Therefore, to determine how this natural acquired humoral immunity to malarial antigens lasts, we determined antigen specific sero-conversion by comparing antigen specific sero-status (Figure 4A) during Day0 and Day14, Day0 and Day365, Day0 and Day730 as indicators of short and long-term antigen specific immunity respectively. Test serum exhibited three different dynamics towards these antigens, some serum samples did not change their serostatus while others changed their serostatus from either seropositive to seronegative or from seronegative to seropositive after 14, 365 or 730 of follow-up days relative to day 0 (Figure 4A and 4B). This change was directly proportional to the total cumulative seropositivity as observed for the protein tertiles (Figure 3B). Proteins in the 1st tertile changed the most; with a mean percent sero-positivity of 43.45% (range 17.19-81.25%), 46.55% (range 18.25-82.48%), 38.25% (range 11.76-75.47%), 32.83% (range 12.12-74.19) on day0, day14, day365 and day730 respectively, and with a mean seroconversion of 3.10% (range −8.00-21.50), −5.20% (range 6.7-7), −10.62% (range −31.25-15.625) from day0 to day14, day0 to day365 and day0 to day730 respectively (supplementary table 2). While proteins in the 2nd tertile changed moderately; with a mean percent sero-positivity of 7.62% (range 0.00-20.31%), 7.58% (range 3.00-17.83), 7.20% (range 0.00-21.15%), 7.04% (range 0.00-35.48%) on day0, day14, day365 and day730 respectively, and with a mean sero-conversion of − 0.037%(range −5.09-5.31%), −0.42%(range −6.37-7.09), −0.57%(range −18.75-24.54%) from day0 to day14, day0 to day365 and day0 to day730 respectively (supplementary table 2). Proteins in the 3^rd^ tertile changed the least; with a mean percent sero-positivity of 0.56% (range 0.00-3.13%), 0.775194%(range 0.00-3.10%), 1.92%(range 0.00-6.00), 2.23%(range 0.00-9.38%) on day0, day14, day365 and day730 respectively, and with a mean sero-conversion of 0.59%(range −1.38-2.94%), 1.36%(range −1.56-6.00%), 1.67%(range −3.13-9.38) from day0 to day14, day0 to day365 and day0 to day730 respectively (supplementary table 2). This shows that humoral immune response against *P. falciparum* antigens change over time in an antigen specific manner that can be generalized into 3 patterns (Figure 4B); immunity to some malaria antigens is acquired quickly and maintained (high percent sero-positivity on day0, day14, day365 and day730 with low or no sero-reversion during day0-day14, day0-day365, day0-day730), or is acquired slowly and not maintained (low percent sero-positivity on day0, day14, day365 and day730 with sero-reversion during day0-day14, day0-day365, day0-day730) or is acquired moderately and maintained or lost (moderate percent sero-positivity on day0, day14, day365 and day730 with moderate or no sero-reversion during day0-day14, day0-day365, day0-day730) (Figure 4B). Antigens that quickly induced immunity included 8MSP4,AMA1, 5MSP3, 606800, 1EBA140, 5SERA9, 1MSP1, P113, PF210C, P41, 13MSP9, 3MSP11, 10MSP6, 2EBA175. Among these, immunity to 8MSP4, AMA1, 1EBA140, 1MSP1, P113, PF210C, P41, 13MSP9, 3MSP11, 10MSP6, 2EBA175 was lost to varying degrees while immunity towards 5MSP3, 606800, 5SERA9 was mostly maintained.

**Figure 4.**
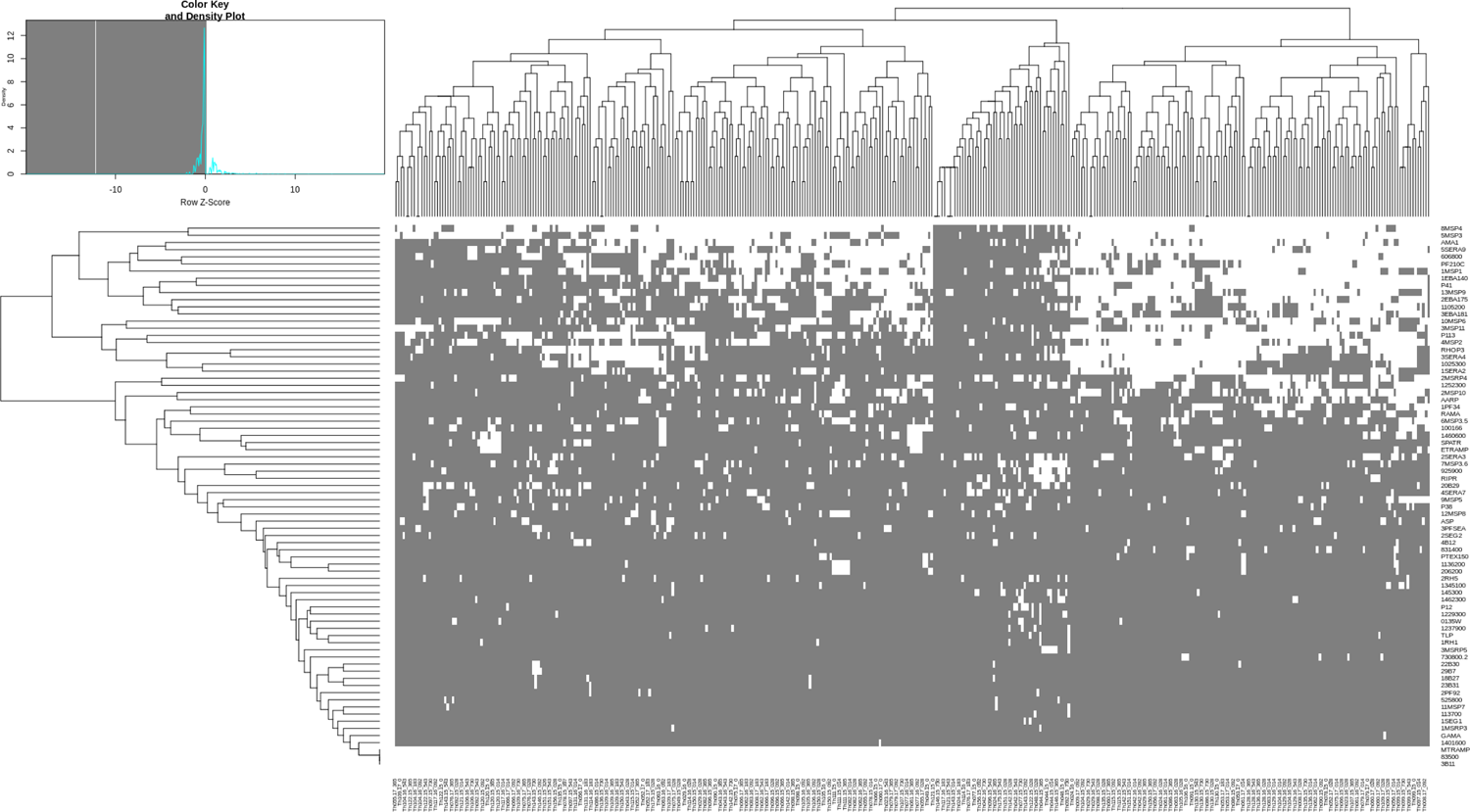
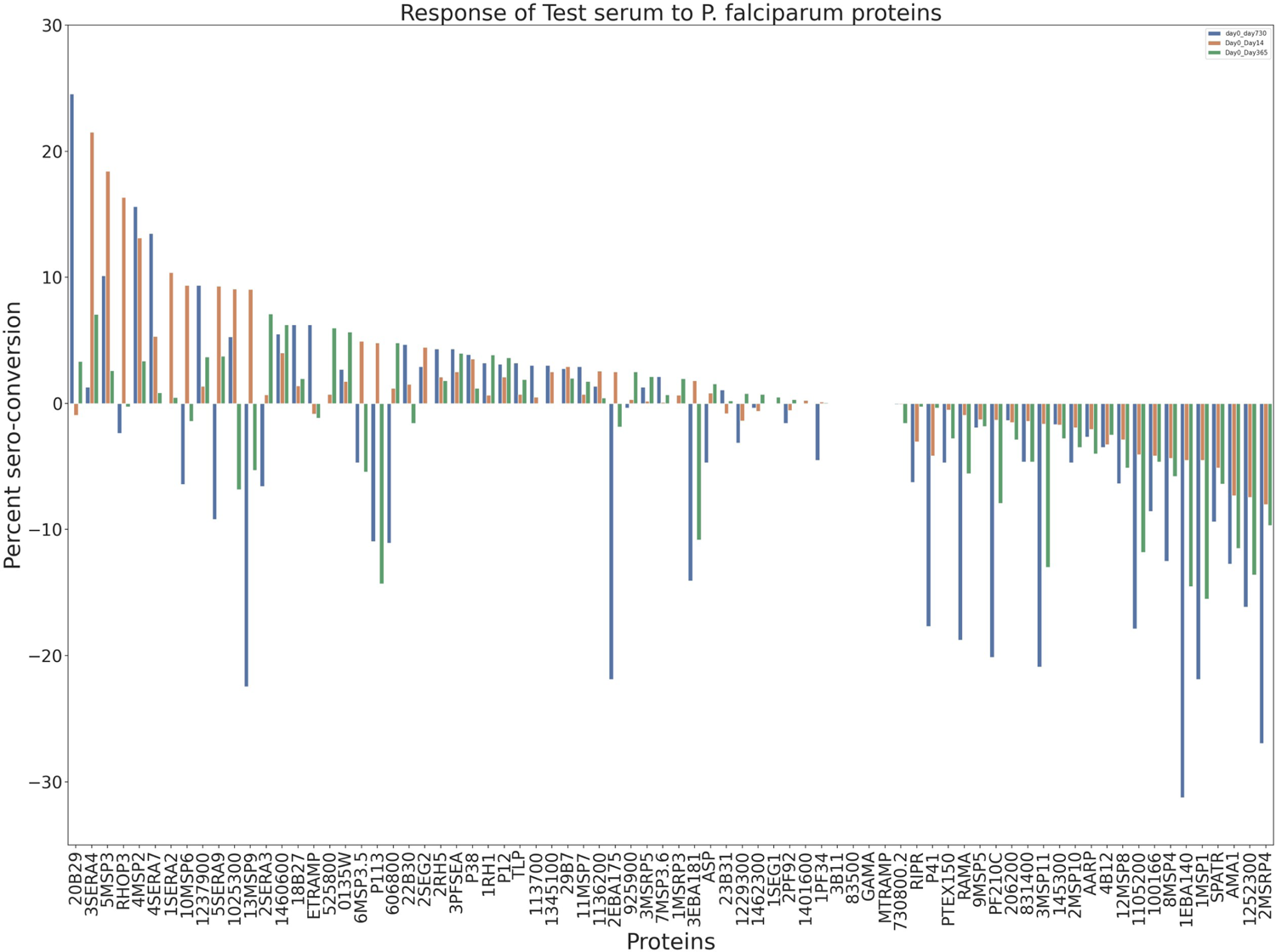
Dynamics in the acquisition and maintenance or loss of humoral immunity towards full length ectodomains *P. falciparum* antigens over time. **A). Antibody acquisition to *P. falciparum* antigens.** A heat-map of sero-status of patient serum from the day of contact (horizontal axis) clustered using hierarchical clustering. White represents sero-positive while gray represents sero-negative. MFI of each test protein (vertical axis) as recognized by test serum per each contact day was divided by mean MFI plus 2 standard deviations of non-immune control serum. Samples with values above 1 were considered as sero-positive while samples with values below 1 were considered as sero-negative. **B). Cumulative prevalence of acquired antibody against *P. falciparum* antigens is antigen specific** A bar plot of sero-conversion (Y-axis) which represents the number of test serums that either maintained their sero-status (no change) or seroreverted (from sero-positive to sero-negative: negative bar-plot) or seroconverted (from sero-negative to sero-positive: positive bar-plot) per each test protein (X-axis) after 14 (blue), 365 (orange) or 730 (green) of follow-up days relative to day 0, as indicated by the color coding.

## DISCUSSION

Using a low transmission setting with few or no malaria re-exposure we set out to study natural immune response in the absence of immune exposure by determining how long immunity to various malarial antigen takes to develop and get lost. Our main goal was to reduce the complications resulting from the influence of potential prevailing immunity from past malaria exposure in a high malaria transmission setting that limit the study of antigen specific dynamics. Furthermore, because cell free protein expression systems, such as *E. coli* and wheat germ are likely to result in incorrect folding and post-translation modification of proteins expressed this way and if coupled with the usage of protein constructs covering protein fragments as opposed to whole proteins, these approaches are highly likely to miss other immunodominant epitopes. All together, these limitations could therefore interfere with epitope integrity, antigen presentation and consequently their recognition by the host immune system. In this study we set out to over-come these limitations by using serum samples obtained from a longitudinal cohort in low malaria transmission setting to perform a high-throughput measurement of natural acquired immunity against *P. falciparum* by utilizing the well established Kilchip micro-array plat-form (Kamuyu et al. 2018), which is a multiplex large panel of mostly full length protein ectodomains (about 70% in total) of novel and known malaria vaccine candidates that were selected based on the *P. falciparum* merozoite surface localised, secreted and invasion proteome (Crosnier et al. 2013; Zenonos, Rayner, and Wright 2014), and then expressed mainly using the mammalian protein expression system. Acquisition of humoral immunity to these full length ectodomains *P. falciparum* antigens demonstrate different dynamics over-time, falling into three categories of the reponse: Highly reactive, moderately reactive and least reactive.

The most reactive antigens inducing high levels of antibodies towards them included 1105200, MSRP4, MSP3.5, 1252300, SERA2, 1025300, EBA140, EBA175, EBA181, MSP3, PF210C, AARP, P113, AMA1, SERA4, MSP2, RHOP3, 606800, SERA9, MSP1, MSP4, MSP10, MSP11, P41, MSP6 and MSP9. There was no sero-reversion towards SERA2, MSP3, SERA4, MSP2, 606800 and SERA9, but only by day 730 for 606800 and SERA9, suggesting that immunity to these antigens is acquired quickly and maintained. Indeed, SERA4 (Zenonos, Rayner, and Wright 2014), 606800 (Osier et al. 2014), SERA2 (Zenonos, Rayner, and Wright 2014), SERA9(Bustamante et al. 2017; Zenonos, Rayner, and Wright 2014), MSP2 (Osier et al. 2014; Richards et al. 2013) and MSP3 (Osier et al. 2014; Sirima et al. 2016) have all been associated with strong protection against malaria. However, MSP3 (Mills et al. 2002), SERA2 (Miller et al. 2002), SERA4 (McCoubrie et al. 2007) and SERA9 (Bustamante et al. 2013) are dispensable during blood-stage. There was sero-reversion towards MSP3.5, 1025300, EBA175, EBA181, P113, RHOP3, MSP6 and MSP9 after day 14 suggesting that immunity to these antigens is acquired quickly and lost slowly. For AMA1, EBA140, 1252300, 1105200, MSRP4, PF210C, AARP, MSP1, MSP4, MSP10, MSP11 and P41 sero-reversion occured by day 14 suggesting that immunity to these antigens is acquired quickly and lost quickly. While RHOP3 is associated with strong protection against malaria (Osier et al. 2014), MSP1, MSP4, MSP10, MSP6 and MSP9 (Osier et al. 2014; Richards et al. 2013) have all been associated with weak protection against malaria. The rest have a mixed association with protection against malaria, especially the erythrocyte binding antigens (EBAs): Moderate to strong association: EBA140 RIII-V, AMA1 and P113 (Osier et al. 2014; Richards et al. 2013). Weak to moderate: EBA175 RIII-V EBA175 RII, EBA181 RIII-V (Osier et al. 2014; Richards et al. 2013). Weak to strong: MSP11 and P41 (Osier et al. 2014; Richards et al. 2013). Infact P41 is also dispensable during blood-stage (Taechalertpaisarn et al. 2012). AARP has low invasion inhibitory activity as part of a Triple Chimeric Antigen of *Plasmodium falciparum* AARP, MSP-3 11 and MSP-1 19 (Kalra et al. 2016).

Moderately reactive antigens inducing moderate levels of antibodies towards them included 145300, 1460600, 831400, PFSEA, 100166, RIPR, MSP8, RH5, SPATR, P38, 1136200, PF34, PTEX150, MSP5, ETRAMP, ASP, RAMA, 206200, MSP3.6, 925900, B29, SERA7 and SERA3. Among these antigens RH5, PFSEA, 1460600, P38, 1136200, SERA7, SERA3, MSP3.6, PF34, ASP, 925900 and B29 had no sero-reversion towards them, but only by day 730 for SERA3, MSP3.6, PF34, ASP, 925900 and day 14 for B29, suggesting that immunity to these antigens is acquired moderately and maintained. Sero-reversion occurred by day 14 towards 145300, 831400, 100166, RIPR, MSP8, SPATR, PTEX150, MSP5, ETRAMP, RAMA and 206200 suggesting that immunity to these antigens is acquired moderately and lost quickly. While 1136200 is strongly correlated with protection against malaria (Osier et al. 2014), P38, RH5 (Osier et al. 2014; Richards et al. 2013) and RIPR (Richards et al. 2013; Bustamante et al. 2013; Reddy et al. 2015) are all weakly to moderately correlated with protection against malaria. PF34 (Osier et al. 2014; Richards et al. 2013), PTEX150, MSP5, ETRAMP, ASP (Osier et al. 2014) and RAMA (Reddy et al. 2015) are all weakly correlated with protection against malaria. PFSEA is associated with protection to *P. falciparum* infection, antibodies against it inhibit egress of schizonts (Raj et al. 2014). 100166 is immuno-reactive and its antibodies inhibit merozoite invasion into erythrocytes (Ito et al. 2019). MSP8, SERA7, SERA3 (Zenonos, Rayner, and Wright 2014) and SPATR (Crosnier et al. 2013), are immuno-reactive towards serum from malaria infected individual. However, SERA7, SERA3 (Miller et al. 2002) and 100166 (Knuepfer et al. 2014) are all dispensable during blood-stage.

The least reactive antigens inducing the lowest levels of antibodies towards them included 1229300, 1462300, 1237900, 730800.2, 113700, 525800, 0135W, MSRP5, MSRP3, RH1, MTRAMP, PF92, MSP7, TLP, P12, GAMA, B30, 1401600, 1345100, 83500, B31 and B27. Among these antigens, 1237900, 113700, 525800, 0135W, MSRP5, MSRP3, RH1, MTRAMP, MSP7, TLP, P12, GAMA, 1401600, 1345100, 83500, B27 and B31 had no sero-reversion towards them suggesting that immunity to these antigens is acquired slowly and maintained at low levels while for 1462300, 730800.2, PF92 and B30 sero-reversion occurred by day 14 suggesting that immunity to these antigens is acquired slowly and lost quickly. While 1229300 is strongly associated with protection against malaria infection (Venkatesh et al. 2019) most of these antigens indeed are weakly correlated with protection against malaria: MTRAMP, PF92, TLP (Osier et al. 2014), MSP7, GAMA, P12 (Osier et al. 2014; Richards et al. 2013) and MSRP3 (Osier et al. 2014; Crosnier et al. 2013). Infact P12 (Taechalertpaisarn et al. 2012), RH1 (Triglia et al. 2004) and 1401600 (Knuepfer et al. 2014) are actually dispensable during blood-stage. 0135W is immunogenic but lowly recognised (Ntumngia et al. 2004). MSRP5 is immunoreactive towards serum from malaria infected individuals (Zenonos, Rayner, and Wright 2014).

Among the above three categories of these antigens, some don’t have longitudinal/serological data and association about them with malaria protection is not known but some have been demonstrated to perform or have been implicated with certain biological processes. These are: 1025300 present in a complex of multiple apicoplast proteins (Mallari et al. 2014), 83500, its synthetic peptides have been shown to inhibit merozoite invasion suggesting that it functions in merozoite invasion (Curtidor et al. 2006), B27/B29/B30/B31, bind to the RBC membrane skeleton, important for the translocation from Maurer’s clefts to the iRBC surface (Zhu et al. 2017), 206200, is a pantothenate transporter and it is essential in parasite development (Augagneur et al. 2013), 925900, is essential for hemozoin formation (Joachim M. Matz et al. 2020; Joachim Michael Matz and Matuschewski 2018), 1345100, is an auxiliary subunit of PTEX (the Plasmodium translocon of exported proteins), it functions in protein export (Egea 2020; Peng, Cascio, and Egea 2015), 1105200, is associated with mRNA-binding in trophozoite and schizont-stage parasites as well as detergent-resistant membrane microdomains in trophozoite blood stage (Bunnik et al. 2016; Yam et al. 2013), MSRP4, it is predicted to be involved in invasion (Ngwa et al. 2017), 1460600, is apart of the IMC sub-compartment proteins (ISPs) together with ISP1 and it is also associated with cytoskeleton in *P. falciparum* female gametocyte (Miao et al. 2017; Poulin et al. 2013; Wang et al. 2020), 145300, is associated with Cytoskeleton in *P. falciparum* female gametocyte (Miao et al. 2017), 1237900, it’s transcription changes in absence of SEMP1 (Dietz et al. 2014), 525800, it’s *P. vivax* ortholog (PVP01_1008000), is among the highly abundant proteins identified in *P. vivax* salivary gland sporozoites (Swearingen et al. 2017) and 1462300, associates with PfRab1 bodies likely being translocated to rhoptires (Morse et al. 2016). Others have not been demonstrated to be involved in or implicated with specific biological processes, these are: MSP3.5, 1252300, 831400, 113700 and 730800.2. Out of these MSP3.5, 1252300, 831400 were among the most reactive proteins, making them high priority candidates ideal for further functional investigation to understand their role in merozoite biology.

Malaria blood stage vaccine candidates that have been studied before individually or in combination with other proteins as part of pre/clinical vaccine trials, such as MSP2 (Lawrence et al. 2000; Genton et al. 2002, 2003), PFSEA (Raj et al. 2014), MSP1 (Ogutu et al. 2009; Chitnis et al. 2015; Sheehy et al. 2011; 2012), MSP3 (Bélard et al. 2011; Sirima et al. 2016), AMA1 (Sheehy et al. 2011; Thera et al. 2011; Srinivasan et al. 2017; Duncan et al. 2011; Ellis et al. 2012; Biswas et al. 2014), EBA175 (Koram et al. 2016; El Sahly et al. 2010) and RH5 (Payne et al. 2017; Douglas et al. 2015) all induced higher levels of antibodies, generally falling under the moderate to most reactive antigens indicating that they all were able to induce high levels of humoral immunity. This acquired immunity changed differently: There was high sero-reversion by day 14 towards EBA175, AMA1 and MSP1 indicating that their long-term immunity was lost quickly while there was no sero-reversion towards RH5, PFSEA, MSP2 and MSP3 indicating that their immunity is long-lasting. While this trend is not similar to other malaria endemic areas, there is variation in the time it takes for antimalaria antibody to decay. While generally levels of antibodies to malaria antigens decline within 6 months (Crompton et al. 2010), antibodies to PfAMA1, PfMSP2, PfMSP1 and EBA175 have a half-life of 10-50 days in the Gambia (White et al. 2014; Akpogheneta et al. 2008), in Ghana there half-life is 10-20 days (White et al. 2014) while in Kenya their half-life is reported to be 10-20 days (Kinyanjui et al. 2007) and >20<=100 years (Ondigo et al. 2014). Classical well known and studied antigens such as MSPs and AMA1 have been moderately to highly correlated with protection against malaria and demonstrated to be highly immune-reactive (Osier et al. 2014; Boyle et al. 2014; Richards et al. 2013; Sirima et al. 2016) but generated strain-specific immunity (Sirima et al. 2016; Ogutu et al. 2009; Chitnis et al. 2015; Sheehy et al. 2011; 2012; Bélard et al. 2011; Thera et al. 2011; Srinivasan et al. 2017; Duncan et al. 2011; Ellis et al. 2012; Biswas et al. 2014; Koram et al. 2016; El Sahly et al. 2010; Ouattara et al. 2013; Genton et al. 2002, 2003; Thera et al. 2011; Barry et al. 2009; E. et al. 2012; Takala et al. 2009; Barry and Arnott 2014; Bailey et al. 2020). On the contrary, novel antigens like RH5, which is currently a promising blood-stage malaria vaccine candidate antigen in Phase 1b clinical trials (Payne et al. 2017; Douglas et al. 2015, 5), has been shown to rarely induce significant levels of anti-PfRh5 antibodies by natural infection, however those produced do associate with protection to varying degrees *in vitro* or in clinical cohorts (Patel et al. 2013; Tran et al. 2014; Richards et al. 2013; Bustamante et al. 2013; Reddy et al. 2015; Douglas et al. 2015) while we here demonstrated that it induces moderate levels of antibodies and immunity against it last-long suggests that it would be ideal to revise the criteria of choosing blood-stage malaria vaccine candidates. We propose that while a potential future malaria blood-stage vaccine candidate(s) besides being essential, having a conserved genetic diversity and the antibodies it induces should have no sero-reversion, it doesn’t necessarily need to have a strong association with protection to malaria nor to have a high i mmune-reactivity profile.

By utilizing a large panel of full length protein ectodomains of novel and known malaria vaccine candidates selected from the *P. falciparum* merozoite proteome (Crosnier et al. 2013; Zenonos, Rayner, and Wright 2014) and expressed using the mammalian protein expression system (Kamuyu et al. 2018), we have demonstrated that these antigens can be recognized by the humoral immune system inducing acquisition of different levels of antibodies and humoral immunity against them, whereby it was either not acquired or if it was acquired, it was either maintained or lost at different rates. Some of the limitations of this study are: absence of a non-exposed baseline in the cohort, Day 0 starts as they present with malaria; the study focused only on blood stage antigens and only less than 100 of them; the composition of the cohort is not broadly representative of the population structure, it didn’t include individuals less than 5 years or greater than 16 years old. We have never-the-less managed to identify several antigens with ability to induce high levels of antibodies after minimal exposure in a low transmission setting making them ideal candidates to be prioritized for further functional or serological studies. These include novel antigens such as PF3D7_1025300, PF3D7_1105800, PF3D7_1334400, PF3D7_0911300, PF3D7_1252300, PF3D7_1460600, PF3D7_1453100, PF3D7_0831400.

## MATERIAL AND METHODS

### Protein microarray probing

*P. falciparum* full length ectodomains were expressed and printed on micro-array slides as described in (Kamuyu et al. 2018) with a slight modification regarding the number of replicate protein spots per each array. Proteins spots were printed in duplicates per each array as opposed to triplicates as done in (Kamuyu et al. 2018). Antigen detection was carried out as well as described in (Kamuyu et al. 2018). Briefly, before the assay samples were diluted at 1:400 in blocking solution (prepared as 2% BSA in Hepes Buffer Saline with 0.1% Tween 20) and arranged in a 96 well plate format. On the day of the assay, printed micro-array slides were pre-scaned using a microarray scanner (GenePix) before starting the assay with PMT set to 400 and power to 100. Four of these pre-scanned slides were set up in 96 well plate format using slide holders. Using a multichannel, 200 µl of washing buffer one (prepared as 1X Hepes Buffer Saline with 0.1% Tween 20) was added to each well, the plates were then sealed and incubated for 5 min at room temperature while shaking at 300 rpm in a hybridization station. After this incubation period the buffer was discarded the and these washing steps repeated two more times. After these washings with wash buffer one, three more washings were done but using wash buffer two (WB2, prepared as 1X Hepes Buffer Saline). 200 µl of blocking solution per well was then added using a multichannel, the plates were then sealed with a plate sealer, covered with a foil paper and incubated for 2 hrs at room temperature while shaking at 300 rpm. The slides were then again washed three times as above with wash buffer one followed with washing buffer two. Diluted samples span at 4000 RPM for 2 minutes s at 4 degrees Celsius, were added at 250 µl of sample per well. The plates were then sealed with a plate sealer and covered with foil paper then incubated overnight at +4 degrees Celsius while shaking at 300 rpm. The slides were then again washed three times as above with wash buffer one followed with washing buffer two. Secondary antibody, diluted at 1:800 in blocking buffer was done using a multichannel, at 150 µl per well. The plates were then sealed with a plate sealer and covered with foil paper then incubated for 3 hours at room temperature while shaking at 300 rpm. The slides were then again washed three times as above with wash buffer one followed with washing buffer two. After these washing steps, the plates were dismantled and then while working on one slide at a time, the slides were dipped several times in distilled water in a Giemsa tray. Eventually, these slides were dry span by putting them upright on a cassette then centrifuging at 1000 rpm for 2 mins at room temperature scanned using a microarray scanner (GenePix) with PMT set to 400 and power to 100.

## Data processing and analysis

Data preprocessing was carried out as described in (Kamuyu et al. 2018). Briefly, data quality check was performed using R v3.6.1 and included evaluation of background intensities, buffer signals, replicate correlation, negative and positive controls. The median fluorescent intensities (MFI) of the local spot background surrounding each spot was subtracted from the MFI of each antigen spot. The extent of within sample variation was then analysed using coefficient of variation (CV). To account for technical slide-to-slide, batch-to-batch variation and mean-variance dependence (MVD) a two-step normalisation process was applied. First, to handle batch-to-batch variation we utilize a wrapper to *SVA’s* function *ComBat()*. Secondly, data normalization was done using variance stabilization normalization (VSN).

## Supplementary information

Please use the any of the following links to access the supplementary information (Supplementary table 1, Supplementary table 2 and Supplementary figure 1A-C): https://datadryad.org/stash/share/xiZ10dUp6j4C4MKrLWxJlFJmw4BtFmBt_720l3k27mQ or https://doi.org/10.5061/dryad.kh189327h

**Supplementary figure 1.** Line plots of percent sero-positive test serums per each contact day over the whole study period against representative test proteins. **A.** Analysis was done including re-infection samples. **B.** Analysis was done excluding re-infection samples. **C.** Analysis was done including samples both with re-infections (blue) and without re-infections (orange) but plotted differently.

## ACKNOWLEDGEMENT

Amy Bei was supported by K01 TW010496. Faith Osier was supported by MRC/DFID African Research Leader Award jointly funded by the UK Medical Research Council (MRC) and the UK Department for International Development (DFID) under the MRC/DFID Concordat agreement (MR/L00450X/1); EDCTP Senior Fellowship (TMA 2015 SF1001); Sofja Kovalevskaja Award from the Alexander von Humboldt Foundation (3.2 - 1184811 - KEN - SKP).

